# Stably bound adaptor proteins modulate directionality of RNP transport

**DOI:** 10.1101/2022.04.12.487826

**Authors:** Ly Jane Phea, Anne Ephrussi

## Abstract

Kinesin-1 and cytoplasmic dynein are molecular motors that mediate long range transport of cargoes along the microtubule cytoskeleton. *oskar* RNA has been documented to switch between the motors during its localization in the *Drosophila* germline syncytium. *oskar* RNA undergoes dynein-mediated transport from the transcriptionally active nurse cells into the oocyte, following which the RNA translocates via kinesin to the posterior pole. Adaptor proteins link the RNA to its motors: the Egalitarian-Bicaudal-D complex links dynein to *oskar* RNA for the initial phase of transport, whereas atypical Tropomyosin 1 (*a*Tm1) links kinesin-1 to *oskar* RNA for the latter phase. Components of the Exon Junction Complex (EJC) as well as the SOLE, a stem loop formed upon splicing of *oskar* RNA, have also been found to be necessary for kinesin-mediated transport of *oskar* RNA. In this study, to dissect the minimal elements required for kinesin-based transport, we tethered *a*Tm1 or kinesin-1 to *oskar* RNA constructs lacking the SOLE. Our results suggest that stably bound *a*Tm1 can indeed bypass the SOLE and EJC to mediate kinesin-1 activity, but the effects of tethered *a*Tm1 are less potent than that of tethered kinesin-1. We also tethered Bicaudal-D to *oskar* RNA, to test whether this would affect kinesin-directed transport of *oskar* RNA, and found that tethered Bicaudal-D directs dynein mediated localization. Our results show that activated Bicaudal-D, along with the recruited dynein, is sufficient for dynein activity. We also show that stable binding of kinesin-1 to the RNA cargo is sufficient for strong kinesin-1 activity. Stably bound *a*Tm1, meanwhile, can only mediate mild kinesin activity, suggesting that other factors may be required to stabilize the binding of kinesin-1 to the RNA cargo.

## Introduction

The intracellular transport of organelles, vesicles, proteins and RNAs to specific locations is central to numerous processes, including establishment of cell polarity and asymmetry during development, responses to signalling, and cell locomotion (Chin & Lécuyer, 2017) and references therein). Long-range transport is especially important in large cells such as the *Drosophila* egg chamber and neurons. Much of the long-range transport in cells occurs along the polarised microtubule cytoskeleton, and the two major classes of microtubule motors involved in transport processes are motor proteins belonging to the kinesin and dynein families. Kinesin-1 usually mediates transport towards the plus ends of microtubules and cytoplasmic dynein towards the minus ends of microtubules.

For RNA transport, adaptor proteins link kinesin-1 and cytoplasmic dynein (hereafter referred to as dynein) to RNA through specific motifs in the RNA (Clark et al., 2007; Dienstbier & Li, 2009; Brendza et al., 2000; Gáspár et al., 2017; Veeranan-Karmegam et al., 2016; Dimitrova-Paternoga et al., 2021). The *Drosophila* egg chamber exists as a syncytium comprising 15 nurse cells and 1 oocyte, surrounded by a monolayer of follicle cells (Spradling, 1993). The oocyte is transcriptionally silent throughout most of oogenesis, and RNAs, proteins and nutrients are transported from the nurse cells into the oocyte by dynein. Among the transported RNAs are maternal RNAs important for the establishment of polarity in the developing *Drosophila* embryo (Barr et al., 2016). The Egalitarian-Bicaudal-D complex links *bicoid*, *gurken* and *oskar* RNAs to dynein for nurse cell to oocyte transport in the egg chamber. The Egalitarian-Bicaudal-D complex also activates dynein for processive movement (Bullock & Ish-Horowicz, 2001; McClintock et al., 2018; McKenney et al., 2014; Clark et al., 2007). Once in the oocyte, *bicoid* and *gurken* RNAs localize in a dynein-dependent manner to the anterior and anterolateral regions of the oocyte respectively, while *oskar* localizes in a kinesin-dependent manner to the posterior pole (Brendza et al., 2000; Ferrandon et al., 1994; Thio et al., 2000; Van De Bor et al., 2005; Ghosh et al., 2012). An atypical isoform of Tropomyosin-1, Tm1-I/C (*a*Tm1), has been identified as an adaptor protein that binds both kinesin-1 and RNA, and is thought to link kinesin-1 to *oskar* RNA in developing egg chambers (Gáspár et al., 2017; Veeranan-Karmegam et al., 2016; Dimitrova-Paternoga et al., 2021).

*In vitro* reconstitution studies have shown that minimal complexes containing dynein or kinesin, an adaptor protein and a cognate RNA cargo are sufficient for cytoskeletal motor-mediated transport (McClintock et al., 2018;Baumann et al., 2020). However, the situation is more complex *in vivo*, as biochemical and live imaging studies suggest that cargo can be simultaneously attached to multiple motor proteins (Kanai et al., 2004; Messitt et al., 2008). Live imaging studies conducted in *Drosophila* egg chambers have also shown that both the velocity and run lengths of RNAs increase when a motor of opposing polarity knocked down or mutated (Mische et al., 2007; Trovisco et al., 2016; Zimyanin et al., 2008). The factors that regulate motor dominance and the eventual directionality of cargo movement in these instances remain poorly understood.

To address how the directionality of transport can be regulated, we focused on *oskar* RNA, which switches from dynein to kinesin-1 based motility during its localization within the germline syncytium (Brendza et al., 2000; Clark et al., 2007; Ephrussi et al., 1991; Jambor et al., 2014; Januschke et al.,2002; Kim-Ha et al., 1991; Palacios & St Johnston, 2002; Sanghavi et al., 2013; Williams et al., 2014;Zimyanin et al., 2008). During the early stages of oogenesis, microtubules nucleate from the oocyte posterior cortex and *oskar* RNA is transported from the nurse cells to the oocyte by dynein (Clark et al., 2007). The oocyte entry signal (OES), a stem loop in the 3’UTR of *oskar* RNA, is necessary for dynein dependent transport of the RNA from the nurse cells into the oocyte (Jambor et al., 2014).

During mid-oogenesis, the oocyte microtubules are re-organized, with plus-ends slightly biased towards the posterior of the oocyte (Jambor et al., 2014; Parton et al., 2011; Cha et al., 2002; Palacios & St Johnston, 2002). *oskar* RNA switches to kinesin transport for its translocation within the oocyte, enriching transiently in the middle of the oocyte before localizing to the posterior pole (V. Zimyanin et al., 2007). Besides *a*Tm1, deposition of the Exon Junction Complex on the *oskar* coding region, as well as the spliced *oskar* localization element (SOLE), a stem-loop structure formed upon *oskar* RNA splicing, were also found to be necessary for kinesin-mediated transport of *oskar* RNA (Hachet & Ephrussi, 2004; Ghosh et al., 2012).

In this study, to dissect the minimal elements required *in vivo* for kinesin-based transport of *oskar* to the posterior of the oocyte, we tethered either *a*Tm1 or kinesin-1 to *oskar* RNA constructs lacking the SOLE. We find that tethering of *a*Tm1 promotes *oskar* RNA localization in the absence of the SOLE and EJC, but that tethered *a*Tm1 is less potent than tethered kinesin-1 in driving the RNA to the posterior pole. In contrast, when we tethered a truncated, constitutively active form of Bicaudal-D to *oskar* RNA, to test whether this would affect kinesin-directed transport of *oskar*, we found that tethered Bicaudal-D localizes *oskar* RNA to the oocyte anterior. Thus, while activated Bicaudal-D can result in strong dynein activity, by recruitment and subsequent activation of the motor, factors other than *a*Tm1 and kinesin-1 are required for strong kinesin-1 activity.

## Results

### Tethered aTm1 promotes *oskar 3’UTR* localization to the posterior of the oocyte

The oocyte entry signal (OES), a stem loop in the 3’UTR of *oskar* RNA, is necessary for dynein dependent transport of the RNA from the nurse cells into the oocyte (Jambor et al., 2014). However, *oskar* RNAs consisting of the 3’UTR (*oskar 3’UTR*) alone or bearing mutations in the SOLE (*oskar-SL1*) fail to localize to the posterior of the oocyte and remain unlocalized, displaying a cloud-like distribution with a slight bias towards the posterior (Hachet & Ephrussi, 2004; Fig 1B). To address whether we could bypass the requirement for the EJC and SOLE and to assess the respective contributions of the adaptors and motors, we engineered MCP-tagged *a*Tm1 to enable tethering to the MS2-tagged *oskar* transcripts (Bertrand et al., 1998; Fig. 1A). To avoid hitchhiking between the *oskar* constructs with endogenous *oskar* RNA, the RNAs were expressed in *oskar* RNA null mutant flies (Jambor et al., 2011). We carried out smFISH to detect the RNAs in the oocyte and used a previously described ImageJ plugin to generate averaged RNA signal distributions and to compute the centre of mass of the RNA distribution, relative to the centre of the oocyte (Gaspar et al.,2014).

**Figure 1:**
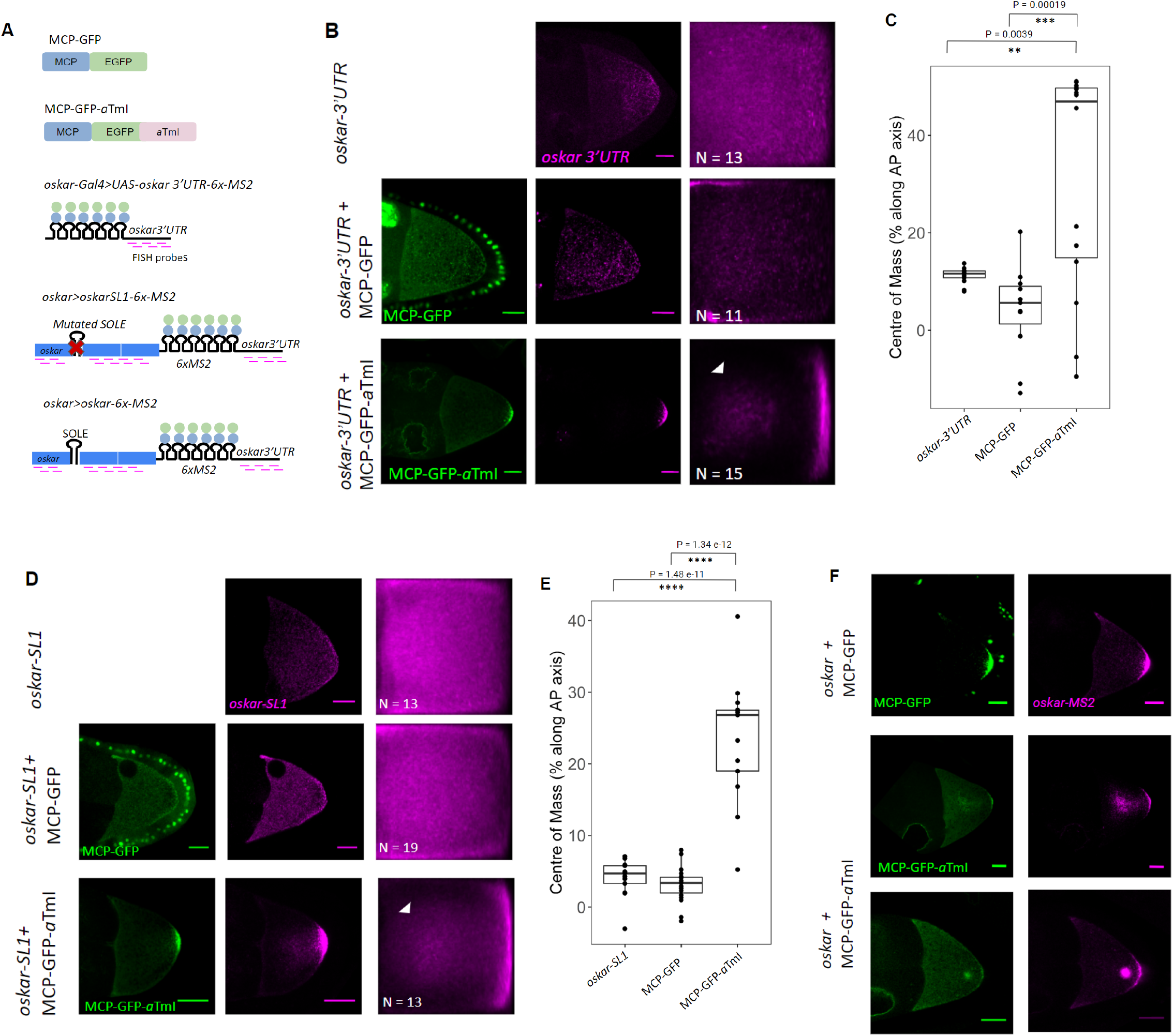
Stably bound *a*Tm1 promotes *oskar 3’UTR* localization to the posterior of the oocyte. (A) Schematic representation of the tethering assay using the MCP-MS2 system, where MCP-GFP or MCP-GFP-*a*TmI were tethered to *oskar 3’UTR, oskar-SL1 or oskar*. (B) Representative images of Stage 9 oocytes expressing *oskar 3’UTR* and MCP-GFP or MCP-GFP-*a*TmI. Panels in green show the GFP signal, whereas the panels in magenta show the RNA signal, detected by smFISH. The right-most panels show the mean distribution of *oskar* RNA signal in Stage 9 oocytes, measured using smFISH; anterior to the left and posterior to the right. N indicates the number of oocytes analysed. White arrowhead indicates central accumulation. Scale bar represents 20 μm. (C) Position of the centre of mass of *oskar 3’UTR* RNA relative to the geometric centre of the oocyte, denoted as 0 in the graph, along the antero-posterior axis. Dots represent individual data points. ANOVA followed by Tukey’s range test were used to compare means. (D) Representative images of Stage 9 oocytes expressing *oskar-SL1* and MCP-GFP or MCP-GFP-*a*TmI. Panels in green show the GFP signal, whereas the panels in magenta show the RNA signal, detected by smFISH. The right-most panels show the mean distribution of *oskar* RNA signal in Stage 9 oocytes, measured using smFISH; anterior to the left and posterior to the right. N indicates the number of oocytes analysed. White arrowhead indicates central accumulation. Scale bar represents 20 μm. (E) Position of the centre of mass of *oskar-SL1* RNA relative to the geometric centre of the oocyte, denoted as 0 in the graph, along the antero-posterior axis. Dots represent individual data points. ANOVA followed by Tukey’s range test were used to compare means. (F) Representative images of Stage 9 oocytes expressing *oskar* and MCP-GFP or MCP-GFP-*a*TmI. Scale bar represents 20 μm.

We found that tethering of *a*Tm1 causes localization of both *oskar 3’UTR* and *oskar-SL1* RNAs to the posterior of the oocyte (Fig. 1B-E). In contrast, tethering of the control MCP-GFP did not promote posterior localization of the RNAs (Fig. 1B-E). Taken together, these results show that tethering *a*Tm1 to localization-impaired *oskar* RNAs can drive their localization to the posterior pole and bypass the requirement for the EJC and the SOLE.

Interestingly, besides their localization to the posterior, tethered *a*Tm1 also led to a central accumulation of the *oskar 3’UTR and oskar-SL1* RNAs (Fig. 1B,D), reminiscent of the central accumulation of *oskar* RNA that normally occurs at stages 7-8 of oogenesis, when microtubules are being reorganized (Cha et al., 2002;Palacios & St Johnston, 2002). To test whether this phenotype might be a result of kinesin hyperactivity, we tethered *a*Tm1 to MS2-tagged wild-type *oskar* RNA. Wild type *oskar* has stronger kinesin activity compared to the non-localizing *oskar* transcripts and thus we expect that any phenotype that arises due to kinesin activity would be stronger in the wild type transcripts. This *oskar-MS2* construct localizes to the posterior pole in a similar manner to endogenous *oskar* RNA, and tethering of MCP-GFP did not affect the localization of *oskar-MS2*.

Upon tethering *a*Tm1, however, we again observed this central localization phenotype, and in some cases, *oskar-MS2* localizes to the posterior with a prominent central dot (Fig. 1F). Thus, our results show that tethered aTm1 leads to the localization of localization-impaired RNAs by increasing the level of kinesin-1 activity. This may occur by direct recruitment of the kinesin-1 motor or by activation of kinesin-1 activity, as kinesin-1 has been shown to exist in an autoinhibited conformation which is only relieved upon cargo binding (Verhey & Hammond, 2009).

### Tethered *a*Tm1 is less effective than tethered kinesin-1 in driving RNA localization

By tethering *a*Tm1 to *oskar 3’UTR* and *oskar-SL1*, we found that *a*Tm1 can drive posterior localization. The central cloud phenotype that we observed also recapitulates a step that occurs during *oskar* RNA localization. To determine whether tethered *a*Tm1 is sufficient for full kinesin-1 activity, whether by recruitment or activation of the motor, we also carried out the experiments using tethered kinesin-1 as a control. As kinesin-1 is required for the establishment of polarity in the *Drosophila* oocyte (Januschke et al., 2002), overexpression of the motor might lead to polarity defects. To circumvent this, we made use of a truncated version of kinesin (Khc-401), which contains only the motor domain and lacks a functional cargo binding domain. Khc-401 was shown to be constitutively active as it lacks autoinhibition, which would also allow us to assess whether full length kinesin-1 requires external activation (Telley et al., 2009).

Expression of either MCP-GFP tagged Khc-401 or full length kinesin-1 (Khc-FL) using a strong driver (*oskar>Gal4*) resulted in formation of egg chambers that did not proceed beyond stage 6 and oogenesis arrest (Fig. S1). This was not observed when Khc-401 and Khc-FL were expressed on their own, in the absence of any *oskar* MS2-constructs. Previous studies have shown that accumulation of *oskar* RNA in the oocyte through dynein-mediated transport is necessary for progression of oogenesis past the early stages (Jenny et al., 2006). Therefore, the termination of oogenesis we observe upon Khc-401 and Khc-FL tethering to *oskar* is most likely due to impairment of nurse cell-to-oocyte transport of *oskar* RNA, resulting in insufficient amounts of *oskar* RNA in the oocyte and oogenesis arrest (Jenny et al., 2006; Kanke et al., 2015). This suggests that the tethering of Khc-401 or Khc-FL to *oskar* RNA interferes with dynein-mediated transport of *oskar* RNA into the oocyte.

To test this and possibly overcome the issue of insufficient *oskar* RNA accumulation in the oocyte and oogenesis arrest, we made use of a weaker Gal4 driver (NGT40) to drive the expression of the MCP fusion proteins. This driver, which is derived from the *nanos* promoter (Tracey et al., 2000), is transcriptionally silent during stages 2-6 of oogenesis. Absence of expression of Khc-401 and Khc-FL during these stages should allow the transport of *oskar* into the oocyte and oogenesis progression.

For both dynein-mediated transport into the oocyte and posterior localization of *oskar* RNA, the effect of kinesin-1 tethering was stronger than that of *a*Tm1 (Fig 2B-E). We observed no significant difference in the behaviour of tethered Khc-FL versus Khc-401. When we tethered *a*Tm1 to *oskar-SL1-MS2*, the amount of *oskar-SL1-MS2* that accumulated in the oocyte was reduced compared to *oskar-SL1-MS2* alone, in the absence of any tethered MCP fusion proteins. The reduction in oocyte accumulation of *oskar* was far more pronounced upon tethering of Khc-401 or Khc-FL, but the effect of tethering the two kinesins was similar (Fig 2B,C). Similarly, when using the weaker driver (NGT40) to drive expression of the MCP construct, we observed that tethered *a*Tm1 drove posterior localization of *oskar-SL1-MS2*, but that tethered kinesin-1 was more effective in localizing the RNA (Fig. 2D,E). Furthermore, the MCP-tagged Khc-FL appeared more concentrated in the posterior domain than did *a*Tm1 or Khc-401 (Fig. 2D). As the levels of expression Khc-FL and Khc-401 were similar (data not shown), we think that Khc-FL may concentrate more efficiently at the posterior. It is notable that this accumulation of Khc-FL at the posterior does not cause a corresponding increase in the amount of *oskar* RNA at the posterior.

**Figure 2:**
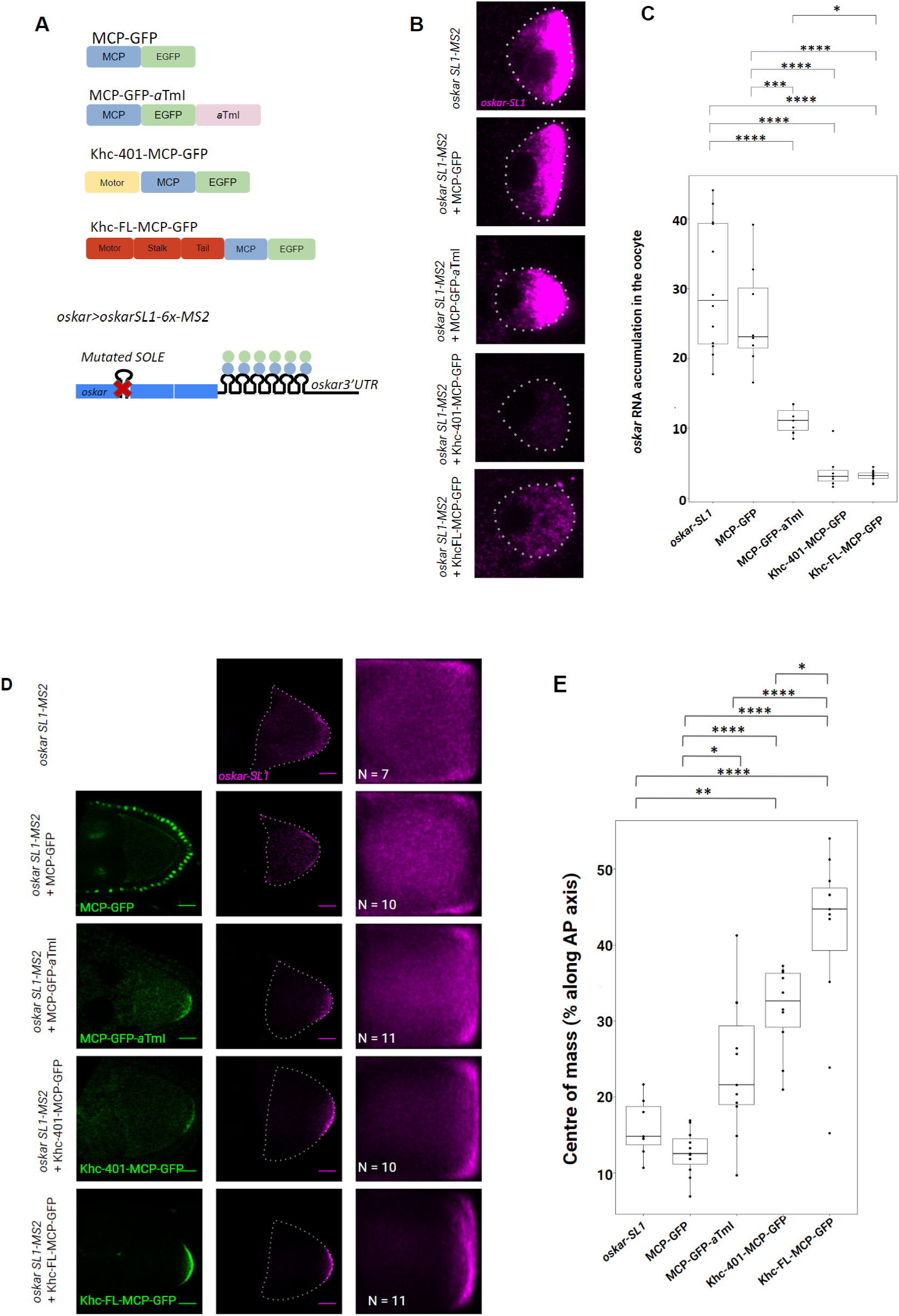
Tethered *a*Tm1 has a weaker effect on localization than tethered kinesin-1. (A) Schematic representation of the tethering assay using the MCP-MS2 system. (B) Representative images of Stage 5 oocytes expressing *oskar-SL1* and MCP-GFP, MCP-GFP-*a*TmI, Khc-401-MCP-GFP or Khc-FL-MCP-GFP. Dotted lines indicate the positions of the oocytes. Scale bar represents 10 μm. (C) Measure of the degree of accumulation of *oskar-SL1* RNA in the oocyte relative to the nurse cells. Dots represent individual data points. ANOVA followed by Tukey’s range test were used to compare means. Significance levels: * <0.1, ** <0.01, **** <0.0001 (D) Representative images of Stage 9 oocytes expressing *oskar-SL1* and MCP-GFP, MCP-GFP-*a*TmI, Khc-401-MCP-GFP or Khc-FL-MCP-GFP. Panels in green show the GFP signal, whereas the panels in magenta show the RNA signal, detected by smFISH. The right-most panels show the mean distribution of *oskar* RNA signal in Stage 9 oocytes, measured using smFISH; anterior to the left and posterior to the right. N indicates the number of oocytes analysed. Dotted lines indicate the position of the oocyte. Scale bar represents 20 μm. (E) Position of the centre of mass of *oskar-SL1* RNA relative to the geometric centre of the oocyte, denoted as 0 in the graph, along the antero-posterior axis. Dots represent individual data points. ANOVA followed by Tukey’s range test were used to compare means. Significance levels: * <0.1, ** <0.01,**** <0.0001

Our results suggest that the stable binding of kinesin-1 to RNA can result in strong kinesin-1 activity in the absence of other activating factors or that the attachment of multiple kinesins can bring about strong kinesin-1 activity.

### Tethered kinesin-1 localizes *bicoid* and *gurken* RNAs to the posterior of the oocyte

Like *oskar, bicoid* and *gurken* RNAs are transported into the oocyte by dynein in early *Drosophila* egg chambers, and the RNAs depend on the dynein machinery to localize to the anterior and anterolateral regions of the oocyte, respectively, during stages 9-10 of oogenesis (Neuman-Silberberg & Schüpbach, 1993; Berleth et al., 1988). To investigate whether adaptor proteins can modulate the directionality of transport, we tethered *a*Tm1 to both transcripts (Fig. 3A). Tethering *a*Tm1 altered the localization of *bicoid* and *gurken*, such that the RNAs spread along the lateral cortex of the oocyte (Fig. 3B, C). In contrast, tethered Khc-401 caused both transcripts to enrich at the oocyte posterior pole (Fig. 3B), albeit only weakly. This is most likely due to the reduced amount of the RNAs accumulating in the oocyte, as we found previously that tethered kinesin-1 interferes with dynein-mediated delivery of RNA from the nurse cell to the oocyte (Fig 2B,C).

**Figure 3:**
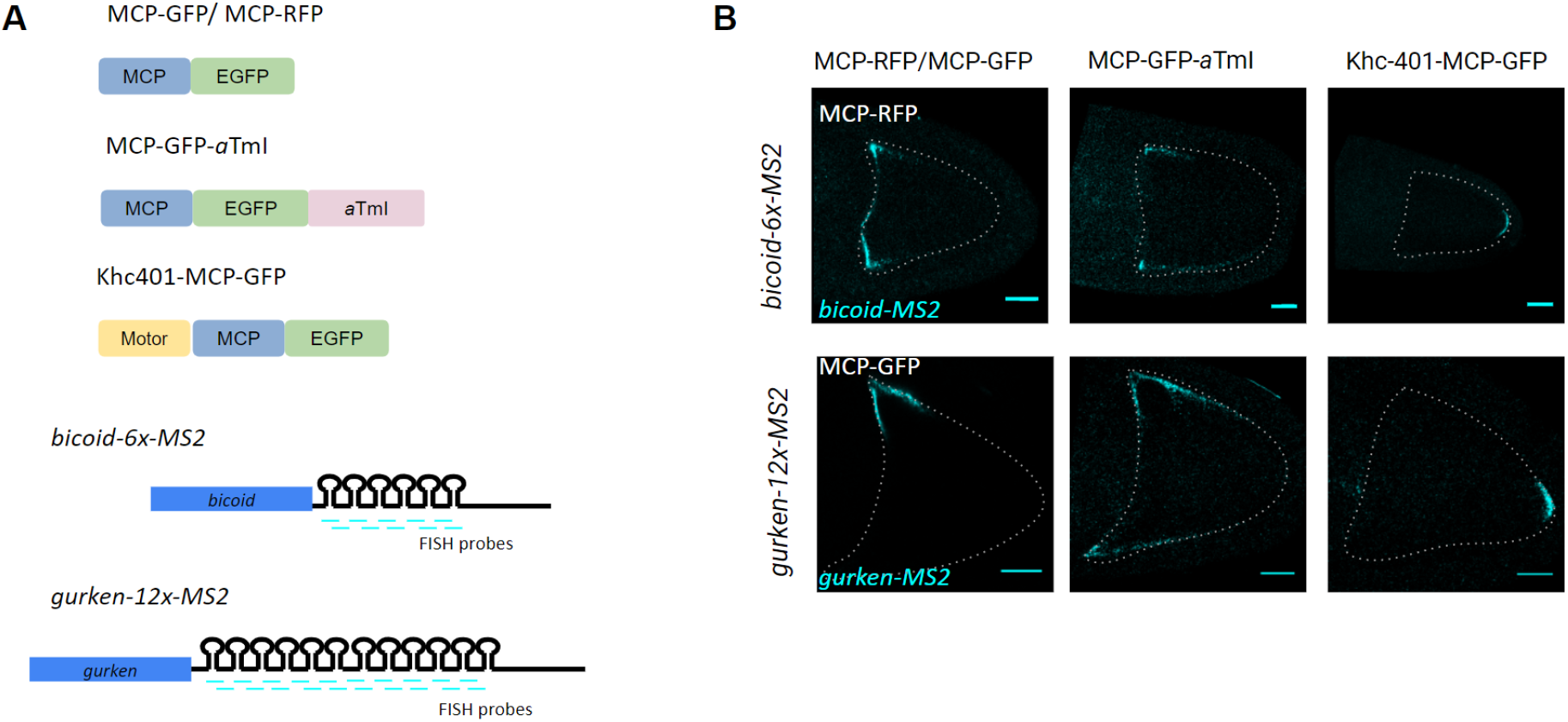
Tethered Khc-401 localizes *bicoid-6x-MS2* and *gurken-12x-MS2* to the posterior of the oocyte. (A) Schematic representation of the tethering assay using the MCP-MS2 system. (B) Representative images of Stage 10 oocytes expressing *bicoid-MS2* or *gurken-MS2* and MCP-GFP/MCP-RFP, MCP-GFP-*a*TmI or Khc-401-MCP-GFP respectively. Panels in cyan show the RNA signal, detected by smFISH. Dotted lines indicate the position of the oocyte. Scale bar represents 20 μm.

Taken together, our results suggest that tethered *a*Tm1, while capable of mediating kinesin-1 activity, is insufficient to establish dominance in a situation where dynein is present and presumably active. The finding that tethered *a*Tm1 is less potent than tethered kinesin-1 also suggests that, *in vivo*, factors other than the adaptor protein are required for strong kinesin-1 activity, which is in line with previous results that point to a requirement of the EJC for kinesin-1 activity (Ghosh et al., 2012).

### Tethered Bicaudal-D-N localizes *oskar-MS2* to the anterior of the oocyte

Having established that *a*Tm1 is capable of mediating kinesin-1 activity, we turned to dynein to investigate whether tethering a dynein adaptor would affect the localization of *oskar* (Fig. 4A). Previous studies have shown that the inclusion of a strong dynein localization signal, such as the K10 TLS, causes ectopic localization of *oskar* RNA at the anterior, and similar phenotypes were observed upon overexpression or dominant-effect mutations of Bicaudal-D (Mach & Lehmann, 1997; Serano & Cohen, 1995). As mentioned previously, RNA cargoes are linked to dynein by the Egalitarian-Bicaudal-D complex. We used a truncated form of Bicaudal-D that contains only the N-terminal coiled-coil (Bicaudal-D-N), as this form of Bicaudal-D mimics the cargo-bound form, wherein autoinhibition is relieved (Dienstbier & Li, 2009). Studies performed *in vitro* have shown that this form of Bicaudal-D promotes processive dynein motility (McKenney et al., 2014; Schlager et al.,2014).

**Figure 4:**
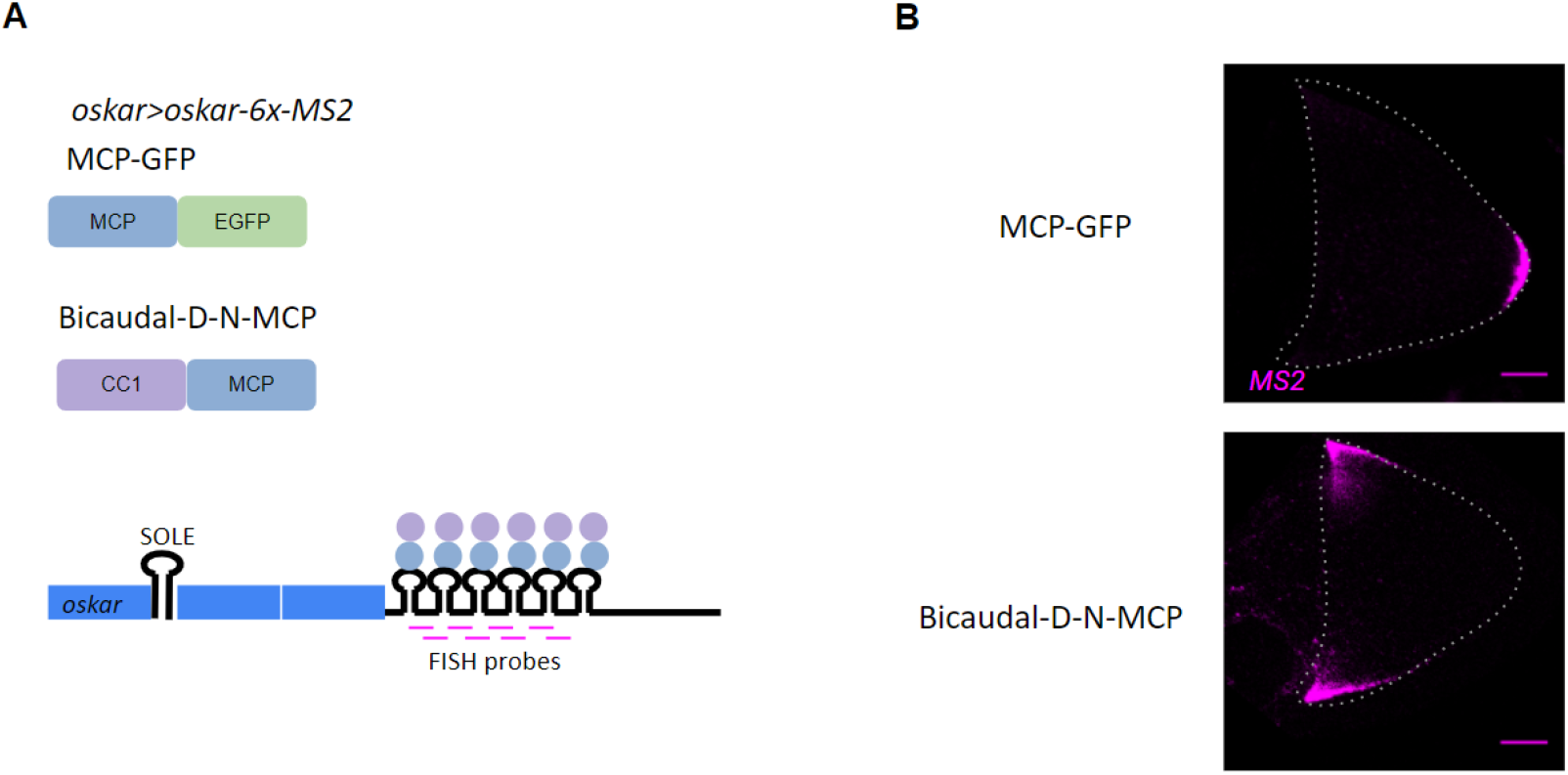
Tethered Bicaudal-D-N-MCP localizes *oskar* to the anterior of the oocyte. (A) Schematic representation of the tethering assay, where MCP-GFP or Bicaudal-D-N-MCP were tethered to *oskar-6x-MS2*. (B) Representative images of Stage 9 egg chambers expressing *oskar-MS2* and MCP-GFP or Bicaudal-D-N-MCP. Specific smFISH probes were used to detect MS2-tagged *oskar*. Panels in magenta show the RNA signal, detected by smFISH. Scale bar represents 20 μm.

We found that tethered Bicaudal-D-N localizes *oskar-MS2* to the anterior (Fig. 4B). *oskar* RNAs containing tethered Bicaudal-D-N also tend to form large clumps at the anterior regions of the oocyte. A previous study showed that localizing RNAs contain more copies of dynein than non-localizing RNAs (Amrute-Nayak & Bullock, 2012). Thus, the clustering of RNAs induced by Bicaudal-D-N may allow RNAs to localize more efficiently to the anterior by recruiting increased numbers of dynein.

## Discussion

Our results show that activated Bicaudal-D, along with the recruited dynein, is sufficient for dynein activity. We also show that stable binding of kinesin-1 to the RNA cargo is sufficient for strong kinesin-1 activity. Stably bound *a*Tm1, meanwhile, can only mediate mild kinesin activity, as other factors may be required to stabilize the binding of kinesin-1 to the RNA cargo. The fact that tethered Bicaudal-D-N results in a stronger localization phenotype than tethered *a*Tm1 may also reflect distinct availability of the kinesin and dynein motor proteins.

A previous study showed that overexpression of *a*Tm1 can promote mild posterior localization of the otherwise unlocalizing *oskar 3’UTR* (Gáspár et al., 2017). This is in contrast to the stronger localization we observe when tethering *a*Tm1. Coupled with the fact that we see strong posterior localization of *oskar-SL1-MS2* upon kinesin-1 tethering, these results indicate that stable binding of *a*Tm1 or of kinesin-1 itself is crucial for posterior localization. The fact that simple overexpression of aTm1 does not cause posterior localization indicates that other factors are required to physically recruit or to stabilize the binding of *a*Tm1 and/or kinesin-1 on *oskar* RNA. As splicing-dependent deposition of the EJC on *oskar* RNA is essential for its posterior localization, the function of the EJC may be to stabilize the association of *a*Tm1 or kinesin-1 with *oskar* RNA.

The stable binding of *a*Tm1 appears to increase the level of kinesin-1 activity on the RNA cargo. This could occur by direct modulation of the motor’s activity, whereby *a*Tm1 stimulates the activity of kinesin-1 by relieving its autoinhibition, or by increasing the number of kinesin motors recruited to the cargo; *a*Tm1 could also increase kinesin activity by directly interfering with dynein activity. Alternatively, *a*Tm1 could exert its effect by modulating the interaction of the motors with microtubules. Thus, *a*Tm might upregulate kinesin activity by acting as a tether and stabilize the motor on microtubules, or downregulate dynein activity by sterically interfering with the dynein binding to microtubules.

The persistent central accumulation of *oskar* RNA we observe upon *a*Tm1 tethering is also observed in hypomorphic kinesin-1 mutants in which the velocity of kinesin was compromised (Serbus et al., 2005). A prominent central dot of *oskar* RNA is also observed in *par-1* mutants, in which oocyte polarity is disrupted (Tomancak et al., 2000;Shulman et al., 2000). In *klar* mutants, where the activity of kinesin is thought to be unrestrained, *oskar* appears to detach from the posterior pole (Gaspar et al., 2014). In *kinesin* mutants in which the region containing the autoinhibitory IAK motif was deleted, Staufen protein, a proxy for *oskar*, localizes in a strong posterior crescent, but is also present in a faint cloud near the posterior region. A previous study also observed this central dot phenotype upon *oskar* overexpression. This phenotype was thought to arise due to the ectopic translation of Oskar protein, which in turn recruits Par-1 kinase, resulting in the stabilization or recruitment of plus ends of microtubules (Zimyanin et al., 2007). In the case of *oskar 3’UTR* and *oskar-SL1*, where Oskar is not translated, we observed a central accumulation of the RNA, however not as a prominent, central dot. This suggests that the prominent central dot phenotype may be dependent on ectopic Oskar production, whereas the diffuse cloud phenotype may occur when kinesin activity is compromised. However, it is unclear whether the diffuse cloud reflects kinesin hyperactivity, or rather, insufficient kinesin activity.

The premature termination of oogenesis we observe upon kinesin-1 tethering to *oskar* RNA suggests that the tethered motor may interfere with dynein-mediated delivery of *oskar* RNA to the oocyte. Indeed, similar phenotypes were observed in mutants of components of the dynein complex, where the activity of dynein is compromised (Ran et al., 1994; Suter & Steward, 1991). It is noteworthy that we did not observe a significant difference in the behaviour of *oskar* RNA whether we tethered Khc-FL or Khc-401. Previous studies identified an autoinhibitory IAK motif in kinesin-1 that is thought to restrict the motor’s activity (Hackney & Stock, 2000; Kaan et al., 2011), however, the removal of this domain did not perturb *oskar* localization (Williams et al., 2014). Although the hinge 2 domain of kinesin-1 is thought to be required for kinesin-1 autoinhibition by allowing the N-terminal motor domain to fold over and interact with the C-terminal tail domain (Moua et al., 2011), mutations in this domain also did not appear to abolish posterior localization of *oskar* RNA (Williams et al., 2014). The fact that IAK and hinge2 mutations do not seem to have a negative impact on kinesin-dependent transport in the *Drosophila* germline suggests that other mechanisms may be responsible for the regulation of *Drosophila* kinesin *in vivo*. A caveat of our result where we observe no significant differences in the behaviour of tethered Khc-FL compared to tethered Khc-401 is that the fusion of MCP-GFP to the C-terminus of kinesin may force it to adopt an open and fully active conformation, or that tethering kinesin to the RNA cargo may be sufficient to alleviate kinesin-1 autoinhibition.

The cortical localization of *bicoid* and *gurken* that we observe upon *a*Tm1 tethering is reminiscent of *oskar* localization in *kinesin-1* (Zimyanin et al., 2007) or *ensconsin* (Sung et al., 2008) mutants. Thus, it may be indicative of no net motor activity, suggesting that tethered *a*Tm1 interferes with dynein activity either directly or by upregulating kinesin-1 activity. Experiments involving injection of *bicoid* RNA into egg chambers also showed that in absence of preincubation with nurse cell cytoplasmic factors, *bicoid* RNA localizes to the lateral cortex of the oocyte (Cha et al., 2001). In this case, bicoid RNA is thought to be unlocalized as it lacks factors required for dynein activity. Thus in the absence of net motor activity, RNAs localize to the cortical regions of the oocyte.

Our tethering experiments show that while stable binding of kinesin-1 to the RNA cargo is sufficient for strong kinesin-1 activity, stably bound *a*Tm1 can only mediate mild kinesin activity. However, our setup does not not allow us to distinguish whether tethered kinesin-1 exerts a stronger effect on the target transcripts by virtue of being tethered or by recruiting endogenous kinesin-1 motors more efficiently than tethered *a*Tm1.

## Materials and Methods

### Generation of transgenic constructs and *Drosophila lines*

oskar 3’UTR-6x-MS2 was amplified from the osk-TRICK plasmid, as described in Halstead et al. (2016) and cloned into the pUASp-attB-ΔK10 vector (Rørth, 1998). The plasmids were injected into VK-33 flies for phiC31 integrase-mediated site-specific integration on chromosome II.

To generate pUASp-attB-ΔK10-TmI, the TmI-I/C ORF and 3’UTR were cloned from the pUASp2-emGFP Tm1RI plasmid (Gáspár et al., 2017) into the BamHI and KpnI sites of the pUASp-attB-ΔK10 vector. MCP (MS2 coat protein) and emGFP sequences were inserted upstream and in frame with the TmI ORF. To generate pUASp-attB-ΔK10-Khc-401-MCP-GFP, the GFP coding sequence was cloned downstream and in frame with the Khc-401-MCP, as described in Gáspár et al. (2017). Similarly, pUASp-attB-ΔK10-KhcFL-MCP-GFP was generated by cloning the Khc ORF in between the KpnI and NotI sites of the pUASp-attB-ΔK10 vector, downstream of the pUAS promoter, and MFP-emGFP sequences were cloned downstream and in frame with the Khc ORF. Plasmids for the MCP-tagged constructs were injected into VK-18 flies for phiC31 integrase-mediated site-specific integration on chromosome II.

*oskar>Gal4VP16* ((Telley et al., 2012); FBtp0083699) and *NGT40* (Tracey et al., 2000);FBti0012287) were used to drive the expression of oskar 3’UTR-6x-MS2 and the MCP-tagged constructs. *oskar-SL1-MS2* and *oskar-WT-MS2* strains were generated by Imre Gáspár and Alessandra Reversi, as described in Ghosh et al. (2012). The expression of *oskar 3’UTR* was driven using the UAS/Gal4 system, as initial attempts to clone it downstream of the endogenous *oskar* promoter did not result in any expression of the transcript.The expression of *oskar-SL1-MS2* was driven from a genomic *oskar* promoter (Ghosh et al., 2012).

Flies expressing the *oskar 3’UTR-MS2* transgene were used in combination with the *oskar-attP-3P3GFP* strain (Gáspár et al., 2017) to generate an *oskar* RNA null background.

The *bicoid-MS2, gurken-MS2* and *Bic-D-N-MCP* fly strains used in this study were kind gifts of Tim Weil, Trudi Schüpbach and Simon Bullock, respectively.

### smFISH

smFISH was performed as described in Gáspár et al. (2017). Probes specific for *oskar, oskar 3’UTR* and *MS2* were designed and labelled with Atto-633 (ATTO-TEC) according to the protocol described in Gáspár et al. (2017). Flies were fed with dry yeast overnight prior to dissection. Ovaries were dissected in PBS and fixed for 20 mins in 2% Paraformaldehyde (Thermo Fisher) in PBST(PBS+0.1% TritonX-100). Fixed ovaries were washed twice in PBST for 10 minutes and incubated with a final probe concentration of 1nM/probe in 250μL HYBEC solution (2x SSC, 15% ethylene carbonate (Sigma Aldrich), 1mM EDTA, 50 μM heparin (Sigma Aldrich), 0.1 mg/mL salmon sperm DNA (Invitrogen), 0.1% Triton X-100) at 37°C. Incubation was carried out for 2 hours for *oskar* probes and 5 hours for *MS2* probes. The ovaries were then washed twice in HYBEC wash solution (2x SSC, 15% ethylene carbonate (Sigma Aldrich), 1mM EDTA, 0.1% Triton X-100) for 15 minutes at 37°C to remove excess smFISH probes. A final wash was performed with PBST for 10 minutes at room temperature, and the ovaries were mounted in Vectashield Antifade mounting medium.

### Microscopy

Imaging was performed on a Leica TCS SP8 confocal microscope using a 63x/1.30 glycerol-immersion objective. GFP signal was imaged using the fixed 488 nm laser line, and signal from the smFISH probes labelled with Atto-633 (ATTO-TEC) was imaged using the fixed 640 nm laser line. Deconvolution of the images were performed using the adaptive Lightning module.

### Analysis of RNA distribution

To quantify the degree of enrichment of RNA signal in the oocytes relative to the nurse cells in early stage egg chambers, the FIJI software was used. The boundaries of the oocytes and nurse cells were delineated manually, and the mean fluorescence of the FISH signal within the oocytes was divided by the mean fluorescence values obtained for the corresponding nurse cells. For background correction, the mean fluorescence values obtained from the follicle cells were subtracted from the values obtained for the nurse cells and for the oocytes.

To obtain a statistically averaged RNA distribution, the CortAnalysis Fiji plugin was used, as described in Gaspar et al. (2014). The outlines of the oocytes were manually defined, the smFISH signal measured and redistributed in a 100×100 matrix. The matrices of multiple oocytes were then projected to give an averaged signal intensity distribution. The centre of mass of the signal distribution relative to the geometric centre of the oocyte was obtained and compared using a pairwise Student’s T-test.

### Statistical analysis

All statistical analyses were performed in R studio (https://www.rstudio.com/) and the graphs were plotted using the ggplot2 library (Wickham 2009).

## AUTHOR CONTRIBUTIONS

Conceptualization, L.J.P. and A.E.; Investigation, L.J.P.; Data Analysis, L.J.P; Writing – Original Draft, L.J.P.; Writing – Review & Editing, L.J.P. and A.E; Supervision, A.E.; Funding Acquisition, A.E.

## ACKNOWLEDGEMENTS

We are grateful to the Bloomington Drosophila Stock Center (BDSC), Imre Gáspár, Simon Bullock, Tim Weil and Frank Wippich for providing fly lines and reagents. We thank Imre Gáspár for helpful advice, and the EMBL Drosophila Injection Service (Alessandra Reversi) for their support. We are grateful to Lucia Cassella, Simone Heber, Mainak Bose and members of the Ephrussi lab for critically reading the manuscript. This work and L.J.P. were supported by EMBL and DFG-FOR 2333 grants EP 37/2-1 and EP 37/4-1 to A.E.

