## Supplemental Figure 1 for "Stably bound adaptor proteins modulate directionality of RNP transport"

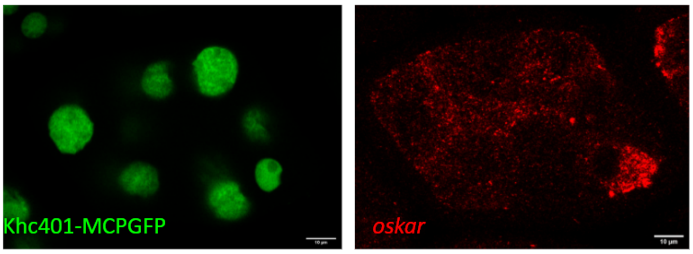


**Figure S1 : Tethered Khc-401-MCP-GFP interferes with oogenesis progression**
Representative images of Stage 5 egg chambers expressing *oskar-SL1* and Khc401-MCP-GFP. Panels in green show the GFP signal, whereas the panels in red show the RNA signal, detected by smFISH. Scale bar represents 10 µm.
